# Resolving microsatellite genotype ambiguity in populations of allopolyploid and diploidized autopolyploid organisms using negative correlations between allelic variables

**DOI:** 10.1101/020610

**Authors:** Lindsay V. Clark, Andrea Drauch Schreier

**Affiliations:** Department of Crop Sciences, University of Illinois, Urbana-Champaign, 1201 W. Gregory Drive, Urbana, IL 61801, USA; Department of Animal Science, University of California – Davis, Davis, CA 95616, USA

**Author notes:** Correspondence: Lindsay V. Clark.

**Keywords:** polyploidy, R package, simple sequence repeat (SSR), sturgeon

## Abstract

A major limitation in the analysis of genetic marker data from polyploid organisms is non-Mendelian segregation, particularly when a single marker yields allelic signals from multiple, independently segregating loci (isoloci). However, with markers such as microsatellites that detect more than two alleles, it is sometimes possible to deduce which alleles belong to which isoloci. Here we describe a novel mathematical property of codominant marker data when it is recoded as binary (presence/absence) allelic variables: under random mating in an infinite population, two allelic variables will be negatively correlated if they belong to the same locus, but uncorrelated if they belong to different loci. We present an algorithm to take advantage of this mathematical property, sorting alleles into isoloci based on correlations, then refining the allele assignments after checking for consistency with individual genotypes. We demonstrate the utility of our method on simulated data, as well as a real microsatellite dataset from a natural population of octoploid white sturgeon (*Acipenser transmontanus*). Our methodology is implemented in the R package polysat version 1.5.

## Introduction

Polyploidy, both recent and ancient, is pervasive throughout the plant kingdom (Udall & Wendel, 2006), and to a lesser extent, the animal kingdom (Gregory & Mable, 2005). However, genetic studies of polyploid organisms face considerable limitations, given that most genetic analyses were designed under the paradigm of diploid Mendelian segregation. In polyploids, molecular markers typically produce signals from all copies of duplicated loci, causing difficulty in the interpretation of marker data (Dufresne *et al*, 2014). If signal (e.g. fluorescence in a SNP assay, or peak height of microsatellite amplicons in capillary electrophoresis) is not precisely proportional to allele copy number, partial heterozygotes may be impossible to distinguish from each other (e.g. AAAB vs. AABB vs. ABBB) (Clark & Jasieniuk, 2011; Dufresne *et al*, 2014). However, under polysomic inheritance (all copies of a locus having equal chances of pairing with each other at meiosis), it is possible to deal with allele copy number ambiguity using an iterative algorithm that estimates allele frequencies, estimates genotype probabilities, and re-estimates allele frequencies until convergence is achieved (De Silva *et al*, 2005; Falush *et al*, 2007). Genotypes cannot be determined with certainty using such methods, but population genetic parameters can be estimated.

The situation is further complicated when not all copies of a locus pair with each other with equal probability at meiosis. “Disomic inheritance” refers to situations in which the locus behaves as multiple independent diploid loci (Obbard *et al*, 2006); similarly, one could refer to an octoploid locus as having “tetrasomic inheritance” if it behaved as two tetrasomic loci. In this manuscript we will refer to duplicated loci that do not pair with each other at meiosis (or pair infrequently) as “isoloci” after Obbard *et al* (2006). When a genetic marker consists of multiple isoloci, it is not appropriate to analyze that marker under the assumption of polysomic inheritance; for example, if allele A can be found at both isoloci but allele B is only found at one isolocus in a population, the genotypes AAAB and AABB are possible but ABBB is not (excluding rare events of meiotic pairing between isoloci). Markers from autopolyploids that have undergone diploidization are likely to behave as multiple isoloci; a locus may still exist in multiple duplicated copies, but the chromosomes on which those copies reside may have diverged so much that they no longer pair at meiosis, or pair with different probabilities (Obbard *et al*, 2006). This segregation pattern is also typically the case in allopolyploids, in which homeologous chromosomes from two different parent species might not pair with each other during meiosis. Further, meiotic pairing in allopolyploids may occur between both homologous and homeologous chromosome pairs, but at different rates based on sequence similarity (Gaeta & Pires, 2010; Obbard *et al*, 2006), which often differs from locus to locus even within a species (Dufresne *et al*, 2014). Waples (1988) proposed a method for estimating allele freqencies in polyploids under disomic inheritance, although it requires that allele dosage can be determined in heterozygotes (in his example, by intensity of allozyme bands on a gel) and allows a maximum of two alleles per locus, with both isoloci posessing both alleles. De Silva *et al* (2005) describe how their method for estimating allele frequencies under polysomic inheritance, allowing for multiple alleles, can be extended to cases of disomic inheritance, but require that isoloci have non-overlapping allele sets, and do not address the issue of how to determine which alleles belong to which isolocus.

Given that marker data do not follow straighforward Mendelian laws in polyploid organisms, they are often recoded as a matrix of ones and zeros reflecting the presence and absence of alleles (sometimes referred to as “allelic phenotypes”; Obbard *et al*, 2006). In mapping populations such binary data can be useful if one parent is heterozygous for a particular allele and the other parent lacks that allele, in which case segregation may follow a 1:1 ratio and can be analyzed under the diploid testcross model (Swaminathan *et al*, 2012; Rousseau-Gueutin *et al*, 2008) (other ratios are possible, in which case the testcross model does not apply). However, in natural populations, inheritance of dominant (presence/absence) markers typically remains ambiguous, and such markers are treated as binary variables that can be used to assess similarity among individuals and populations but are inappropriate for many population genetic analyses, *e.g.* tests that look for departures from or make assumptions of Hardy-Weinberg Equilibrium (Clark & Jasieniuk, 2011).

Microsatellites are a special case given that they have multiple alleles, allowing for the possibility of assigning alleles to isoloci, which would drastically reduce the complexity of interpreting genotypes in allopolyploids and diploidized autopolyploids. For example, if an allotetraploid individual has alleles A, B, and C, and if A and B are known to belong to one isolocus and C to the other, the genotype can be recoded as AB at one isolocus and CC at the other isolocus, and the data can be subsequently processed as if they were diploid. If two isoloci are sufficiently diverged from each other, they may have entirely different sets of alleles. This is in contrast to other markers such as SNPs and AFLPs that only have two alleles (except in rare cases of multi-allelic SNPs), in which case isoloci must share at least one allele (or be monomorphic, and therefore uninformative). With microsatellites, one could hypothetically examine all possible combinations of allele assignments to isoloci and see which combination was most consistent with the genotypes observed in the dataset, but this method would be impractical in terms of computation time and so alternative methods are needed. Catalán *et al* (2006) proposed a method for assigning microsatellite alleles to isoloci based on the inspection of fully homozygous genotypes in natural populations. In their example with an allotetraploid species, any genotype with just two alleles was assumed to be homozygous at both isoloci, and therefore those two alleles could be inferred to belong to different isoloci. With enough unique homozygous genotypes, all alleles could be assigned to one isolocus or the other, and both homozygous and heterozygous genotypes could be resolved. However, their method made the assumption of no null alleles, and would fail if it encountered any homoplasy between isoloci (alleles identical in amplicon size, but belonging to different isoloci). Moreover, in small datasets or datasets with rare alleles, it is likely that some alleles in the dataset will never be encountered in a fully homozygous genotype. The method of Catalán *et al* (2006) was never implemented in any software to the best of our knowledge, despite being the only published methodology for splitting polyploid microsatellite genotypes into diploid isoloci.

In this manuscript, we present a novel methodology for assigning microsatellite alleles to isoloci based on the distribution of alleles among genotypes in the dataset. Our method is appropriate for natural populations, as long as the dataset can be split into reasonably-sized groups of individuals (∼ 100 individuals or more) lacking strong population structure. It is also appropriate for certain mapping populations, including F_2_, recombinant inbred lines, and doubled haploids. It can be used on organisms of any ploidy as long as each subgenome has the same ploidy, for example octoploid species with four diploid subgenomes or two tetraploid subgenomes, but not two diploid subgenomes and one tetraploid subgenome. Negative correlations between allelic variables are used to cluster alleles into putative isolocus groups, which are then checked against individual genotypes. If necessary, alleles are swapped between clusters or declared homoplasious so that the clusters agree with the observed genotypes within a certain error tolerance. Genotypes can then be recoded, with each marker split into two or more isoloci, such that isoloci can then be analyzed as diploid or polysomic markers. Our method works when there are null alleles, homoplasy between isoloci, or occasional meiotic recombination between isoloci, albeit with reduced power to find the correct set of allele assignments. We test our methodology on simulated allotetraploid, allohexaploid, and allo-octoploid (having two tetrasomic genomes) data, and compare its effectiveness to that of the method of Catalán *et al* (2006). We also demonstrate the utility of our method on a real dataset from a natural population of octoploid white sturgeon (*Acipenser transmontanus*). Our methodology, as well as a modified version of the Catalán *et al* (2006) methodology, are implemented in the R package polysat version 1.5.

## Materials and Methods

### Rationale

Say that a microsatellite dataset is recoded as an “allelic phenotype” matrix, such that each row represents one individual, and each allele becomes a column (or an “allelic variable”) of ones and zeros indicating whether that allele is present in that individual or not. Under Hardy-Weinberg equilibrium and in the absence of linkage disequilibrium, these allelic variables are expected to be independent if the alleles belong to different loci or different isoloci. However, if two alleles belong to the same locus (or isolocus), the allelic variables should be negatively correlated. This is somewhat intuitive given that the presence of a given allele means that there are fewer locus copies remaining in which the other allele might appear (Fig. 1A). The negative correlation can also be proved mathematically (Supplementary Materials and Methods). We use “correlation” in a broad sense here; “negative correlation” means that the presence of one allele is associated with the absence of another allele or vice versa.

**Figure 1:**
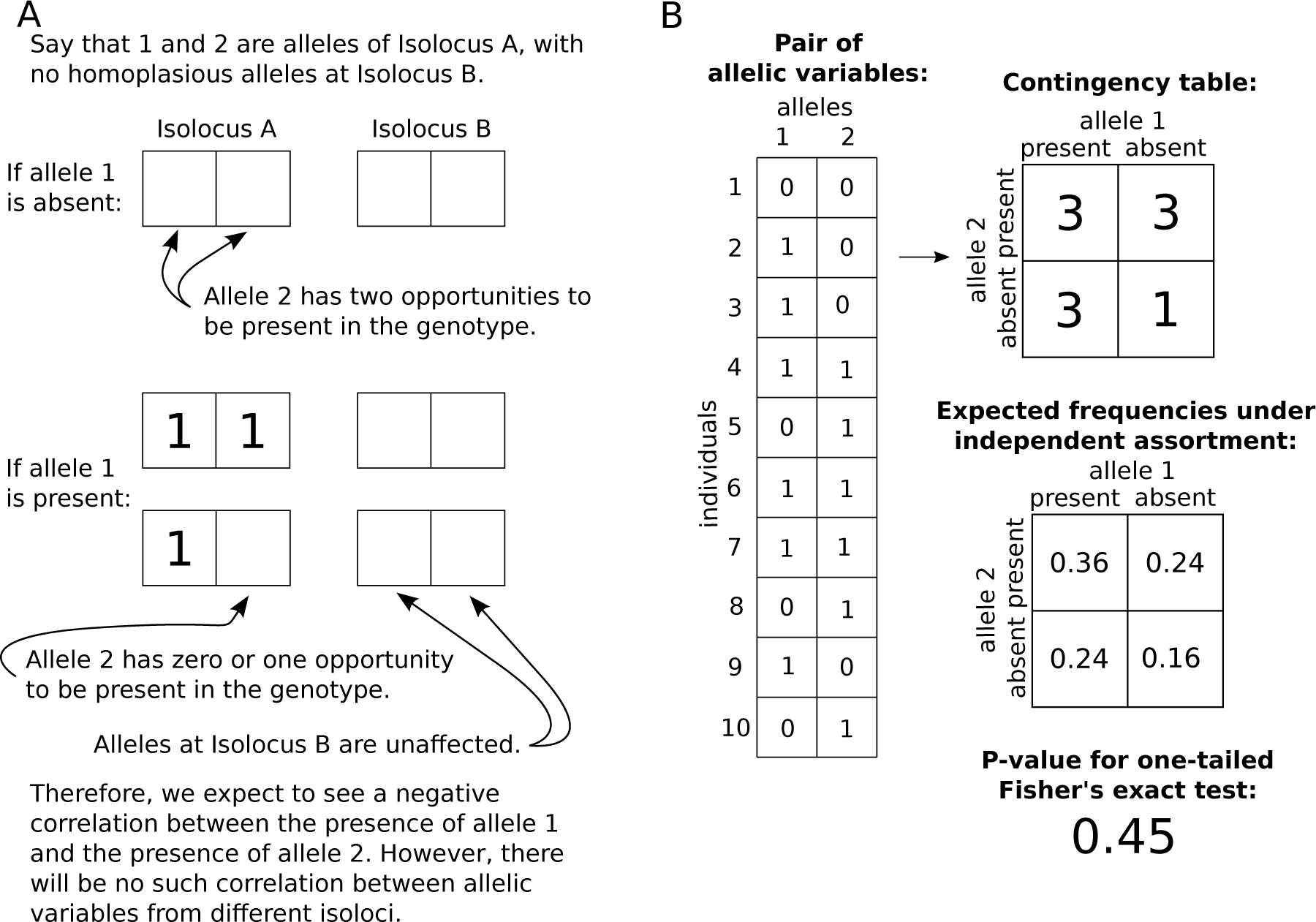
Negative correlation between two allelic variables at a locus. (A) Qualitative reasoning for the expectation of negative correlation between two allelic variables at the same isolocus. (B) Use of Fisher’s exact test to identify negative correlation between a pair of allelic variables. Ten individuals are shown for the sake of illustration, but an ideal dataset would have 100 or more individuals. In the allelic variables, presence of an allele in an individual is indicated by 1, and absence is indicated by 0.

## Algorithm for clustering alleles into isoloci

### Preliminary clusters: the alleleCorrelations function

An overview of our algorithm is presented in Fig. 2. To test independence of two binary allelic variables, we use Fisher’s exact test since it is appropriate for small sample sizes, which are likely to occur in typical population genetics datasets when rare alleles are present. A 2-by-2 contingency table is generated for the test, with rows indicating presence or absence of the first allele, columns indicating presence or absence of the second allele, and each cell indicating the number of individuals in that category (Fig. 1B). A one-tailed Fisher’s exact test is used, with the alternative hypothesis being that more individuals just have one allele of the pair than would be expected if the allelic variables were independent, *i.e.* the alternative hypothesis is that the odds ratio is less than one, indicating a negative association between the presence of the first allele and the presence of the second allele. This alternative hypothesis corresponds to the two alleles belonging to the same isolocus, whereas the null hypothesis is that they belong to different isoloci and therefore assort independently. The P-values from Fisher’s exact test on each pair of allelic variables from a single microsatellite marker are then stored in a symmetric square matrix. We expect to see clusters of alleles with low P-values between them; alleles within a cluster putatively belong to the same isolocus. For clustering algorithms, zeros are inserted along the diagonal of the matrix, since the P-values are used as a dissimilarity statistic. The function `alleleCorrelations` in polysat 1.5 produces such a matrix of P-values for a single microsatellite marker. The same function also produces two sets of preliminary assignments of alleles to isoloci, using UPGMA and the Hartigan & Wong (1979) method of K-means clustering, respectively. The `n.subgen` argument is used to specify how many subgenomes the organism has, *i.e* into how many isoloci each locus should be split.

**Figure 2:**
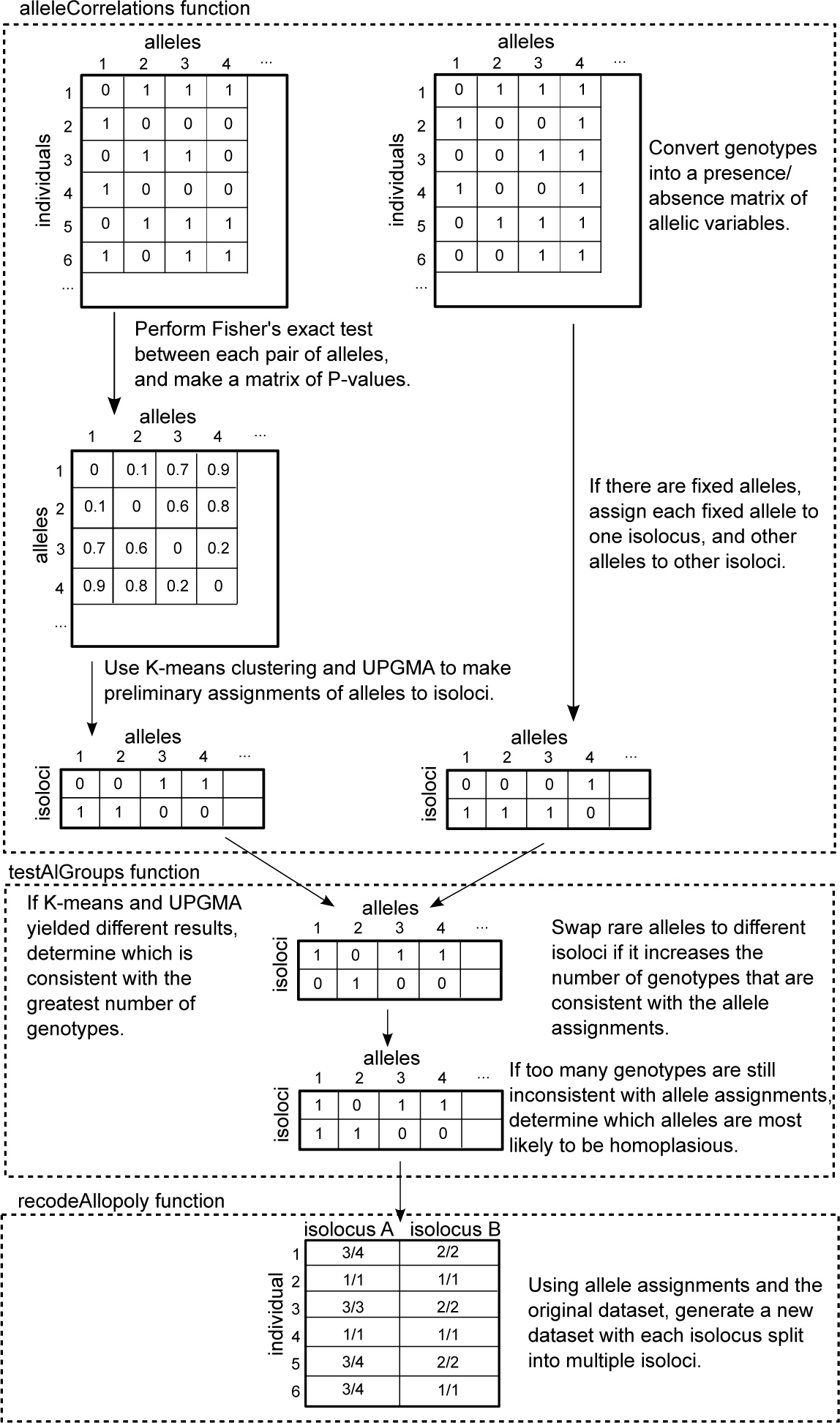
Overview of functions in polysat 1.5 for processing allopolyploid and diploidized autopolyploid datasets. Additionally, the processDatasetAllo function can be used to automatically run alleleCorrelations and testAlGroups on every locus in a dataset. In the box representing the alleleCorrelations function, all alleles belonging to the locus on the left are variable in the dataset, so Fisher’s exact test is used to find correlations between allelic variables, then K-means and UPGMA are used to perform clustering. The locus on the right has one allele (4) that is present in all individuals, making it impossible to assign alleles to isoloci using Fisher’s exact test. In the box representing the testAlGroups function, all steps are performed on all loci regardless of whether or not fixed alleles are present.

Population structure can also cause correlation between allelic variables, for example if two alleles are both common in one subpopulation and rare in another. Because correlation caused by population structure can potentially obscure the correlations that are used by our method, the `alleleCorrelations` function checks for significant positive correlations (after Holm-Bonferroni multiple testing correction) between allelic variables, which could only be caused by population structure, scoring error (such as stutter peaks being mis-called as alleles, and therefore tending to be present in the same genotypes as their corresponding alleles), or linkage disequilibrium (if two isoloci are part of a tandem duplication on the same chromosome, as opposed to duplication resulting from polyploidy), and prints a warning if such correlations are found.

If one or more alleles are present in all genotypes in a dataset, it is not possible to perform Fisher’s exact test to look for correlations between those fixed allelic variables and any others. The function `alleleCorrelations` therefore checks for fixed alleles before performing Fisher’s exact test. Each fixed allele is assigned to its own isolocus. If only one isolocus remains, all remaining alleles are assigned to it. If no isoloci remain (*e.g.* in an allotetraploid with two fixed alleles and several variable alleles), then all remaining alleles are assigned as homoplasious to all isoloci. If multiple isoloci remain (*e.g.* in an allohexaploid with one fixed allele), then Fisher’s exact test, K-means clustering and UPGMA are performed to assign the alleles to the remaining isoloci. It is possible that an allele with a very high frequency may be present in all genotypes but not truly fixed (*i.e.* some genotypes are heterozygous). However, allele swapping performed by `testAlGroups` (see below) can assign alleles to an isolocus even if that isolocus already has an allele assigned to it that is present in all individuals.

### Corrected clusters: the testAlGroups function

Although K-means was more accurate overall than UPGMA using simulated data (Supplementary Table 1), UPGMA sometimes assigned alleles correctly when K-means assigned them incorrectly. To choose between K-means and UPGMA when they give different results, the function `testAlGroups` in polysat checks every genotype in the dataset against both results. Assuming no null alleles or homoplasy (which are dealt with later in the algorithm), a genotype is consistent with a set of allele assignments if it has at least one allele belonging to each isolocus, and no more alleles belonging to each isolocus than the ploidy of that isolocus (*e.g.* two in an allotetraploid). The ploidy of isoloci is specified using the `SGploidy` argument. The set of results that is consistent with the greatest number of genotypes is selected, or K-means in the event of a tie. Selecting the best results out of K-means and UPGMA improved the accuracy of allele assignments at all ploidies, particularly hexaploids (Supplementary Table 1).

**Table 1:** Percentages of simulated datasets with correct allele assignments under different levels of population structure. Two populations of 50 allotetraploid individuals were simulated under different allele frequencies, then merged into one dataset that was then used for making allele assignments. The value shown in the leftmost column was randomly added or subtracted from the frequency of each allele in the first population to generate the allele frequencies of the second population. For isoloci with odd numbers of alleles, one allele had the same frequency in both populations. For each difference in allele frequency, 1000 simulations were performed (5000 total). *F_ST_* was calculated from allele frequencies as (*H_T_* − *H_S_*)/*H_T_*, where *H_S_* is the expected heterozytosity in each subpopulation, averaged across the two subpopulations, and *H_T_* is the expected heterozytosity if the two subpopulations were combined into one population with random mating. Means and standard deviations across 1000 simulations are shown for *F_ST_*. The third column shows the percentages of datasets in which significant positive correlations were detected between any pair of alleles; positive correlations can be used as an indication that there is population structure in the dataset. The fourth and fifth columns indicate the percentages of datasets with correct allele assignments, using our method and that of Catalán *et al* (2006). 95% confidence intervals are given for percentages.

| Difference in allele frequency | $F_{ST}$ | Significant positive correlations | K-means + UPGMA + swap $\leq 0.50$ | Catalán |
| --- | --- | --- | --- | --- |
| 0.0 | $0.000 \pm 0.000$ | $0\% \pm 0\%$ | $94\% \pm 1\%$ | $84\% \pm 2\%$ |
| 0.1 | $0.016 \pm 0.004$ | $2\% \pm 1\%$ | $99\% \pm 1\%$ | $89\% \pm 2\%$ |
| 0.2 | $0.062 \pm 0.013$ | $21\% \pm 3\%$ | $93\% \pm 2\%$ | $94\% \pm 1\%$ |
| 0.3 | $0.117 \pm 0.021$ | $62\% \pm 3\%$ | $88\% \pm 2\%$ | $99\% \pm 1\%$ |
| 0.4 | $0.176 \pm 0.026$ | $82\% \pm 2\%$ | $88\% \pm 2\%$ | $100\% \pm 0\%$ |

We expected that rarer alleles would be more likely to be assigned incorrectly, given that they would be present in fewer genotypes and therefore there would be less statistical power to detect correlations between their variables and other allelic variables. To correct the allele assignments, an algorithm was added to the `testAlGroups` function that individually swaps the assignment of each rare allele to the other isolocus (or isoloci) and then checks whether the new set of assignments is consistent with a greater number of genotypes than the old set of assignments. If an allele is successfully swapped, then every other rare allele is checked once again, until no more swaps are made. The maximum number of genotypes in which an allele must be present to be considered a rare allele is adjusted using the `rare.al.check` argument to the `testAlGroups` function. We found that swapping alleles present in ≤ 50% of genotypes (`rare.al.check = 0.5`) improved the accuracy of the algorithm (Supplementary Table 1), so we used that setting in all evaluations of the algorithm except where noted otherwise. Note that the frequency of genotypes with a given allele will always be higher than the allele frequency itself, although a 50% threshold is still much higher than the cutoff for considering an allele to be “rare” in most population genetic analyses.

Although our algorithm attempts primarily to sort alleles into non-overlapping groups, there is always a possibility that different isoloci have some alleles with identical amplicon sizes (homoplasy). Therefore, we introduced an algorithm to the `testAlGroups` function to check whether any genotypes were still inconsistent with the allele assignments after the allele swapping step, and assign alleles to multiple isoloci until all genotypes (or a particular proportion that can be adjusted with the `threshold` argument, to allow for meiotic or scoring error) are consistent with the allele assignments. The allele that could correct the greatest number of inconsistent genotypes (or in the event of a tie, the one with the lowest P-values from Fisher’s exact test between it and the alleles in the other isolocus) is made homoplasious first, then all genotypes are re-checked and the cycle is repeated until the desired level of agreement between allele assignments and genotypes is met.

Mutations in primer annealing sites are a common occurrence with microsatellite markers, and result in alleles that produce no PCR product, known as null alleles. One potential issue with null alleles is that, when homozygous, they can result in genotypes that do not appear to have any alleles from one isolocus. Such genotypes are used by the `testAlGroups` function as an indicator that alleles should be swapped or made homoplasious, which would be incorrect actions if the genotype resulted from a null allele rather than inaccuracy of allele assignment. We therefore added an argument to the `testAlGroups` function, `null.weight`, to indicate how genotypes with no apparent alleles for one isolocus should be prioritized for determining which alleles to assign as homoplasious. If null alleles are expected to be common, `null.weight` can be set to zero so that genotypes with no apparent alleles for one isolocus are not used for assigning homoplasy. The default value of 0.5 for `null.weight` will cause `testAlGroups` to use genotypes with no apparent alleles for one isolocus as evidence of homoplasy, but with lower priority than genotypes with too many alleles per isolocus. (No argument was added to adjust the allele swapping algorithm, since it only swaps alleles if the overall agreement with the dataset is improved.)

### Recoding datasets based on allele assignments: the processDatasetAllo and recodeAllopoly functions

The function `processDatasetAllo` is a wrapper function that runs `alleleCorrelations` and `testAlGroups` in sequence on every marker in the dataset. It tests several parameter sets for `testAlGroups`. If the dataset was divided into subpopulations to prevent bias from population structure, allele assignments from the same parameter set are merged across subpopulations using the `mergeAlleleAssignments` function. `processDatasetAllo` generates a series of plots to indicate assignment quality, and selects a suggested best parameter set for each locus by first selecting the parameter set that results in the least amount of missing data when the genotypes are recoded, or in the case of a tie the parameter set that results in the fewest homoplasious alleles.

The list of allele assignments (output by `processDatasetAllo`) and the original dataset are then passed to the `recodeAllopoly` function, which produces a new dataset in which each marker is split into multiple isoloci. Missing data are substituted for genotypes that cannot be resolved due to homoplasy in the allele assignments. (For example, if alleles A and B belong to different isoloci, and C belongs to both, the genotype ABC could be AA BC, AC BB, or AC BC, assuming no null alleles.) An argument called `allowAneuploidy` lets the user specify whether to allow for apparent meiotic error. If `allowAneuploidy = TRUE`, for genotypes with too many alleles for one isolocus, the function will adjust the recorded ploidy for the relevant samples and isoloci. (Ploidy is used by other polysat functions, such as those that estimate allele frequency.) Otherwise, missing data are inserted where there are too many alleles per isolocus.

### Implementation of the Catalán method: the catalanAlleles function

polysat 1.5 also includes an implementation of the algorithm of Catalán *et al* (2006). One difference between our implementation and the original is that we allow ploidies higher than tetraploid, *e.g.* in a hexaploid, a genotype with three alleles is assumed to be fully homozygous. Additionally, after fully homozygous genotypes are examined, fully heterozygous genotypes are also examined if necessary for assigning alleles that were not present in any fully homozygous genotypes. The output of `catalanAlleles` can be passed directly to `recodeAllopoly`.

## Simulated datasets

The function `simAllopoly` was added to polysat in order to generate simulated datasets for testing the accuracy of allele assignment methods. It simulates one locus at a time, and allows for adjustment of the number of isoloci, the ploidy of each isolocus, the number of alleles for each isolocus, the number of alleles that are homoplasious between isoloci, the number of null alleles (producing no amplicon), allele frequencies in the population, the meiotic error rate (frequency at which different isoloci pair with each other at meiosis), and the number of individual genotypes to output. By default, alleles from the first isolocus are labeled A1, A2, etc., alleles from the second isolocus labeled B1, B2, etc., and homoplasious alleles labeled H1, H2, etc.

For initial evaluation of clustering methods (Supplementary Table 1), 10,000 simulated markers were generated for 100 individuals each for allotetraploid, allohexaploid, and allo-octoploid (two tetrasomic isoloci) species under Hardy-Weinberg Equilibrium. Although not included in the simulated datasets, note that it is also possible for an octoploid to possess four diploid subgenomes, as in strawberry. Each isolocus had a randomly chosen number of alleles between two and eight, and allele frequencies were generated randomly. A set of allele assignments for one marker was considered to be correct if no alleles were assigned incorrectly.

To evaluate the effect of sample size on assignment accuracy, 1000 additional markers were simulated for populations of 50, 100, 200, 400, and 800 individuals for allotetraploid, allohexaploid, and allo-octoploid species.

To simulate population structure, 5000 simulated markers were generated for two populations of 50 allotetraploid individuals. Allele frequencies differed by five fixed amounts (Table 1) between the two populations, with 1000 markers simulated for each amount.

The effect of homoplasy on allele assignment methods was evaluated by simulating 1000 allotetraploid markers each for sample sizes of 50, 100, 200, 400, and 800, and homoplasious allele frequencies of 0.1, 0.2, 0.3, 0.4, and 0.5.

To evaluate allele assignment when null alleles were present, 5000 markers were simulated for 100 allotetraploid individuals, with 1000 simulated markers at each null allele frequency of 0.1, 0.2, 0.3, 0.4, and 0.5.

Occasional pairing between homeologous (in an allopolyploid) or paralogous (in an autopolyploid) chromosomes may occur during meiosis. As a result, offspring may be aneuploid, having too many or too few chromosomes from either homologous pair, or may have translocations between homeologous or paralogous chromosomes. Most commonly, the aneuploidy or translocations will occur in a compensated manner (Chester *et al*, 2015), meaning that for a given pair of isoloci, the total number of copies will be the same as in a non-aneuploid, but one isolocus will have more copies than expected and the other isolocus will have fewer (e.g. three copies of one isolocus and one copy of the other isolocus in an allotetraploid). To evaluate the accuracy of allele assignment for isoloci that occasionally pair at meiosis, 4000 markers were simulated for 100 allotetraploid individuals, with 1000 simulated markers at each meiotic error rate of 0.01, 0.05, 0.10, and 0.20.

A custom script was written to simulate genotypes in allopolyploid mapping populations. Allotetraploid, allohexaploid, and allo-octoploid (with two tetrasomic subgenomes) populations were simulated, with 200 individuals in each population. For each ploidy, 1000 loci were simulated for each generation spanning F_2_ to F_8_, assuming completely homozygous parents. Allele assignments were performed with the `alleleCorrelations` and `testAlGroups` functions, with `null.weight = 1` and `rare.al.check = 0.25`.

## Empirical dataset

To demonstrate the usefulness of our allele assignment method on a real dataset, we used previously published data from natural populations of octoploid white sturgeon (*Acipenser transmontanus*; Drauch Schreier *et al*, 2012). Previous studies of inheritance patterns in this species suggested that it possesses two tetrasomic subgenomes, at least for portions of its genome (Rodzen & May, 2002; Drauch Schreier *et al*, 2011). We selected for allele assignment the eight microsatellite markers that, based on number of alleles per genotype, appeared to be present in eight copies rather than four.

Because population structure can impact allele clustering, we first performed a preliminary analysis of population structure using the `Lynch.distance` dissimilarity statistic in polysat and principal coordinates analysis (PCoA) using the `cmdscale` function in R. Thirteen microsatellite markers were used for PCoA, including the eight used for allele assignment and five tetrasomic (present in four copies rather than eight) markers. Allele assignment methods were then tested on the whole dataset and on a subpopulation identified by PCoA.

The `testAlGroups` function was run on the sturgeon dataset with and without allele swapping (`rare.al.check` set to 0.5 and 0, respectively). In checking for homoplasy, we allowed up to 5% of genotypes to disagree with allele assignments in anticipation of meiotic error, scoring error, or genotypes homozygous for null alleles (`tolerance = 0.05`), and to allow for null alleles at low frequency we set `null.weight = 0.5` so that genotypes with too many alleles per isolocus would be used for assignment of homoplasy first, before genotypes with no alleles for one of their isoloci.

To evaluate the accuracy and usefulness of allele assignments, we compared *G_ST_* (Nei & Chesser, 1983) estimates using the five tetrasomic loci to estimates using the putatively tetrasomic recoded isoloci. Pairs of isoloci were excluded from *G_ST_* estimates if they had any homoplasious alleles. Allele frequencies for tetrasomic loci and isoloci were estimated using the method of De Silva *et al* (2005) using the `deSilvaFreq` function in polysat with the selfing rate set to 0.0001. Pairwise *G_ST_* between sampling regions was then estimated with the `calcPopDiff` function in polysat.

## Results

### Simulated natural populations

For all ploidies, we found that the accuracy of both our method and the Catalán *et al* (2006) method was dependent on sample size, and that our method performed better than the Catalán *et al* (2006) method at all sample sizes (Fig. 3). For tetraploids and hexaploids, the effect of sample size was greater on the Catalán *et al* (2006) method than on our method, particularly at small sample sizes. For octoploids, the success of the Catalán *et al* (2006) method was near zero even with 800 individuals in the dataset (due to the low probabiltiy of producing fully homozygous genotypes at tetrasomic isoloci), whereas our method had an accuracy of 93% with 800 octoploid individuals.

**Figure 3:**
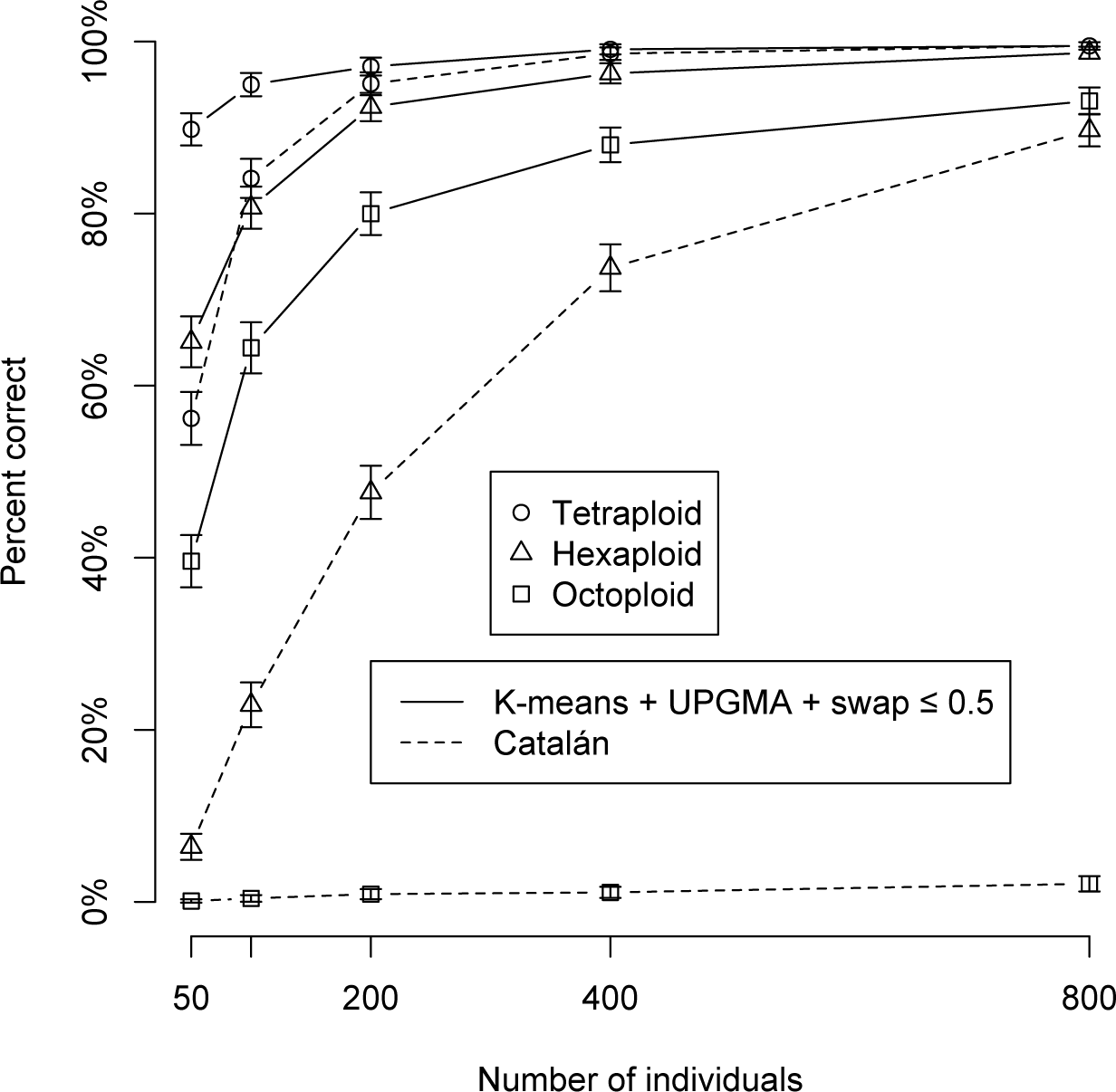
Accuracy of allele assignments with different sample sizes. For each ploidy and sample size, 1000 simulations were performed. Octoploids were simulated with two tetraploid genomes. Whiskers indicate 95% confidence intervals. “Swap ≤ 0:5” indicates that testAlGroups was used with rare.al.check = 0.5. The y-axis indicates the percentage of datasets for which allele assignments were completely correct.

Both negative and positive correlations between allelic variables at different loci can occur when the assumption of random mating is violated by population structure, confounding the use of negative correlations for assigning alleles to isoloci. We found that accuracy of our method remained high (∼ 90%) even at moderate levels of *F_ST_* (∼ 0.2; Table 1). Interestingly, low levels of population structure (*F_ST_* ≈ 0.02) improved the accuracy of our method to 99%, compared to 94% when *F_ST_* = 0 (Table 1), probably as a result of an increase in the number of double homozygous genotypes, which would have been informative during the allele swapping step. For this same reason, the Catalán *et al* (2006) method, which depends on double homozygous genotypes, had an improved success rate as population structure increased, and exceeded our method in accuracy at moderate levels of *F_ST_* (Table 1). However, accuracy of our method decreased with increasing *F_ST_* when *F_ST_* > 0.02 (Table 1), likely because correlations between alleles caused by population structure outweighed the benefits of increased homozygosity. In our simulations, significant postive correlations between allelic variables were found in most datasets that had moderate population structure (Table 1).

One advantage of our method over that of Catalán *et al* (2006) is that our method allows for alleles belonging to different isoloci to have identical amplicon sizes (homoplasy). We tested the accuracy of allele assignments across several sample sizes and frequencies of homoplasious alleles, with and without the allele swapping algorithm (Fig. 4). Allele assignments were most accurate when allele swapping was not performed before testing for homoplasious alleles, and when the homoplasious allele was at a frequency of 0.3 in both isoloci. When allele assignments were correct, we tested the mean proportion of genotypes that were resolvable, given several frequencies of a homoplasious allele (Table 2). Although accuracy of assignment had been highest with a homoplasious allele frequency of 0.3, only 57% of genotypes could be resolved in such datasets (Table 2).

**Figure 4:**
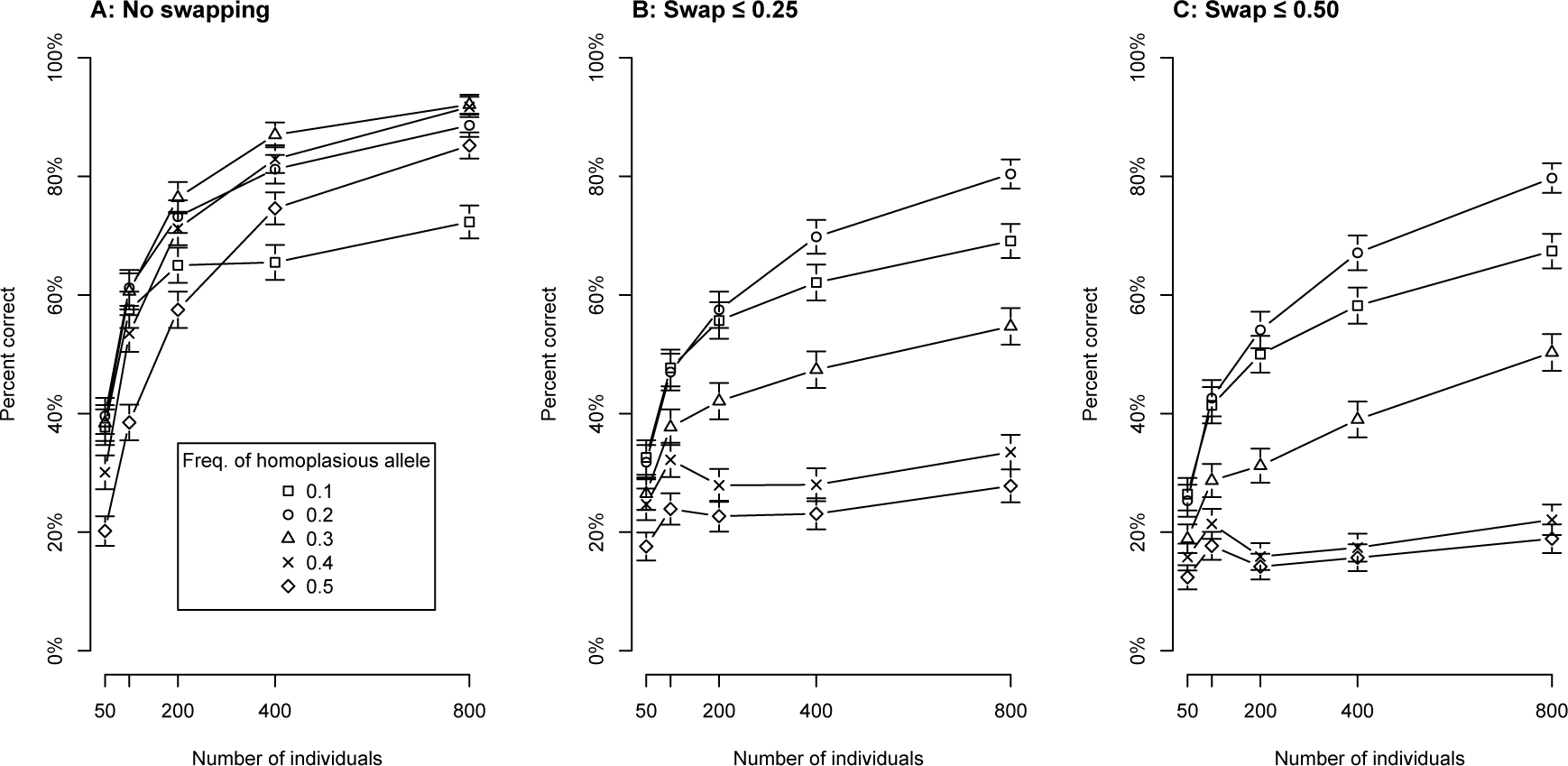
Percentages of simulated datasets with correct allele assignments when homoplasious alleles are present. Whiskers indicate 95% confidence intervals. The y-axis indicates the percentage of datasets for which allele assignments were completely correct. Allotetraploid datasets were simulated with one pair of homoplasious alleles (alleles from two different isoloci, but with identical amplicon size) for each locus. The frequency of homoplasious alleles was identical at both isoloci in each dataset, and was set at five different levels (0.1 through 0.5). Five different sample sizes were tested (50, 100, 200, 400, and 800). For each homoplasious allele frequency and sample size, 1000 datasets were simulated. Allele assignments were made using three methods: K-means + UPGMA (A; rare.al.check = 0), K-means + UPGMA + swap ≤ 0.25 (B; rare.al.check = 0.25), or K-means + UPGMA + swap ≤ 0.50 (C; rare.al.check = 0.5); plus an algorithm in the function testAlGroups that identifies the alleles most likely to be homoplasious, and assigns alleles as homoplasious until all genotypes are consistent with allele assignments.

**Table 2:** For datasets from Fig. 4 with correct allele assignments at `rare.al.check = 0` (no swapping), percentages of genotypes that could be unambiguously resolved. Means and standard deviations are shown.

| Freq. of homoplasious allele | Mean percentage of genotypes that could be resolved |
| --- | --- |
| 0.1 | $85.6\% \pm 5.6\%$ |
| 0.2 | $71.2\% \pm 8.3\%$ |
| 0.3 | $59.4\% \pm 9.4\%$ |
| 0.4 | $51.5\% \pm 9.0\%$ |
| 0.5 | $48.5\% \pm 7.0\%$ |

To test the effect of null alleles on the accuracy of our allele assignment method, we simulated datasets in which one isolocus had a null allele (Fig. 5). We found that, when null alleles were present, the accuracy of the algorithm was greatly improved when genotypes lacking alleles for one isolocus were not used as evidence of homoplasy. We also found that the allele swapping algorithm improved the accuracy of allele assignments when the null allele was at a frequency of 0.1 in the population. However, at higher null allele frequencies (≥ 0.4) allele assignments were more accurate without allele swapping.

**Figure 5:**
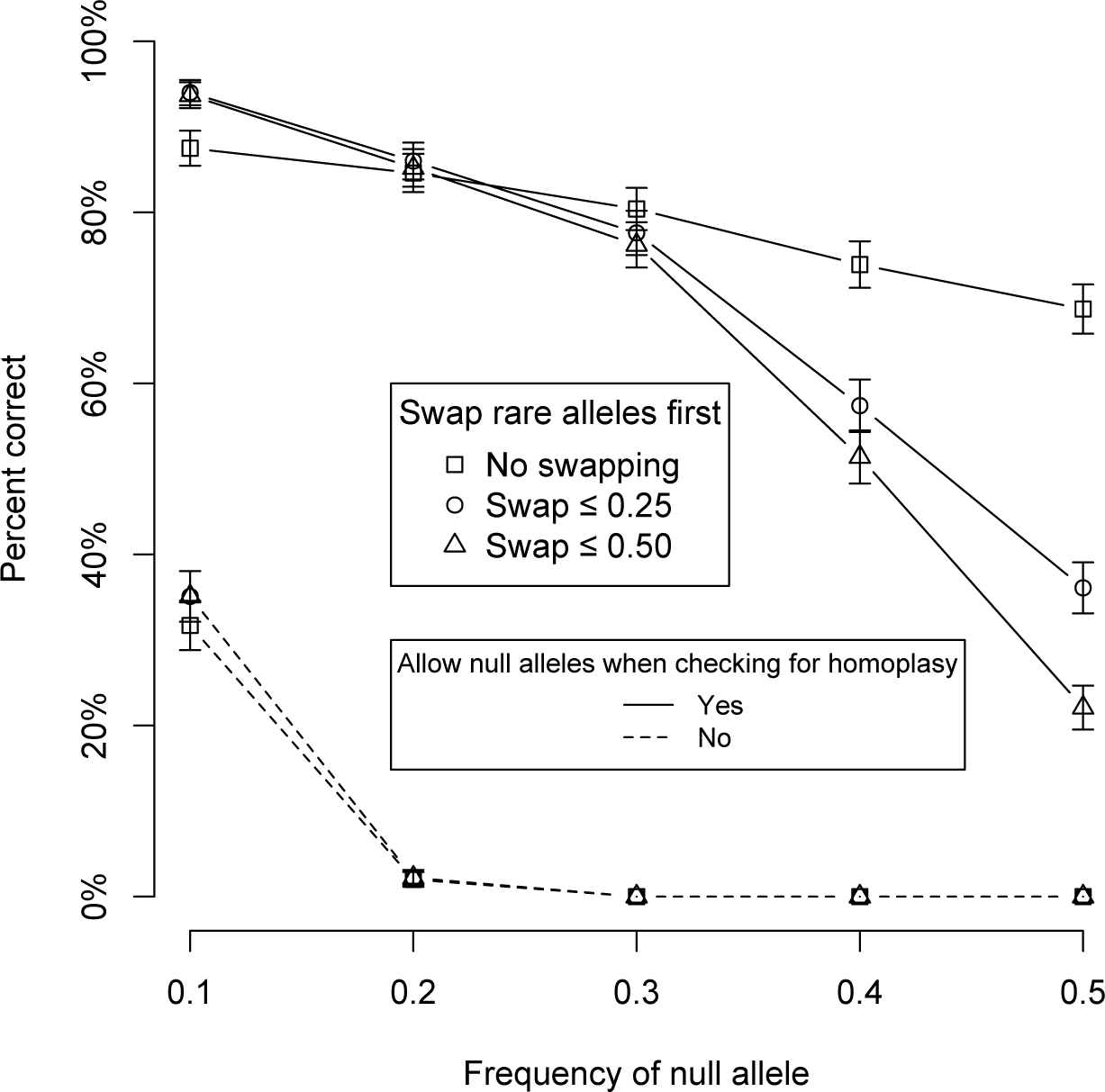
Percentages of simulated datasets with correct allele assignments when one isolocus has a null allele. Whiskers indicate 95% confidence intervals. The y-axis indicates the percentage of datasets for which allele assignments were completely correct. Allotetraploid datasets were simulated, and frequency of the null allele was set at one of five levels (x-axis). 1000 datasets were simulated at each null allele frequency. Two parameters for testAlGroups were adjusted: rare.al.check at values of zero, 0.25, and 0.5 (corresponding to the methods K-means + UPGMA, K-means + UPGMA + swap ≤ 0.25, and K-means + UPGMA + swap ≤ 0.50, respectively); and null.weight at values of zero (null alleles are allowed when checking for evidence of homoplasy) and 0.5 (genotypes lacking alleles belonging to a given isolocus are taken as evidence that their other alleles are homoplasious).

We simulated datasets in which gametes resulting in compensated aneuploidy (meiotic error) occured at a range of frequencies from 0.01 to 0.2 (Fig. 6). At all meiotic error rates, the allele swapping algorithm from `testAlGroups` improved the accuracy of allele assignment (Fig. 6). Meiotic error did not have a large impact on the success of our method; even at a meiotic error rate of 0.2 (where 0.5 would be fully autopolyploid), our algorithm still had an accuracy of 62% on datasets of 100 individuals with no homoplasy, null alleles, or population structure (Fig. 6).

**Figure 6:**
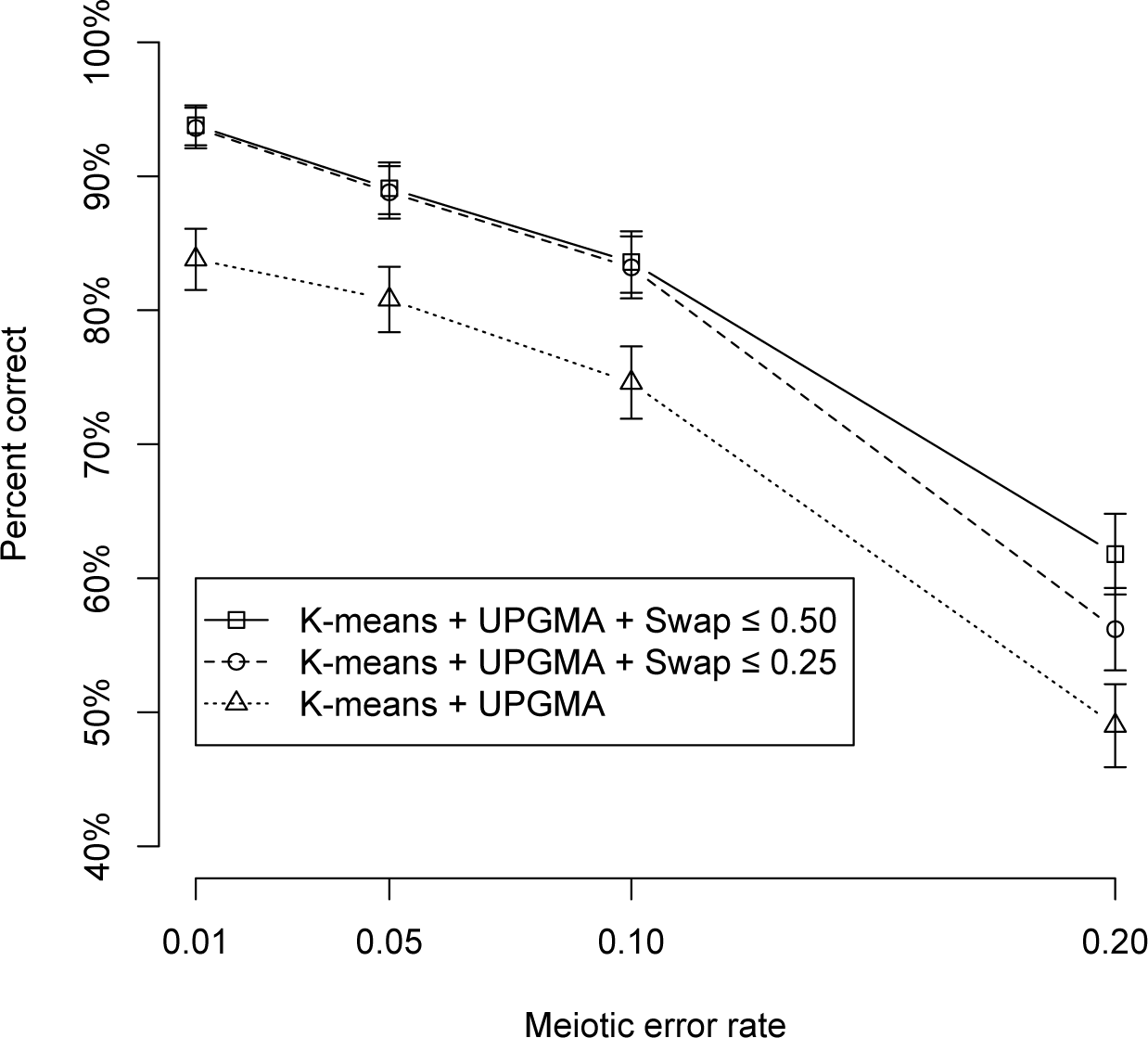
Percentages of simulated datasets with correct allele assignments when meiotic error causes compensated aneuploidy. Whiskers indicate 95% confidence intervals. The y-axis indicates the percentage of datasets for which allele assignments were completely correct. Meiotic error was simulated in the simAllopoly function on a per-gamete basis, with each error causing an allele from one isolocus to be substituted with an allele from the other isolocus. Each dataset was otherwise simulated for an allotetraploid organism with 100 individuals. Meiotic error rate, as shown in the x-axis, was controlled using the meiotic.error.rate argument of simAllopoly. For each error rate, 1000 datasets were simulated. For the testAlGroups function, the tolerance argument was set to 1 to prevent the function from checking for homoplasy, and rare.al.check was set to zero, 0.25, or 0.5 (corresponding to the methods K-means + UPGMA, K-means + UPGMA + swap ≤ 0.25, and K-means + UPGMA + swap ≤ 0.50, respectively). Each dataset was tested for all three values of rare.al.check.

We also examined the effect of number of alleles on the accuracy of our method. Accuracy was highest when the number of alleles was similar among isoloci (Supplementary Table 2).

## Assignment of alleles to isoloci in octoploid sturgeon

When using principal coordinates analysis to test for genetic structure prior to performing allele assignment, we identified two major genetic groups (Supplementary Table 3, Supplementary Fig. 1) that were similar to the population structure previously observed (Drauch Schreier *et al*, 2012). The smaller group (Pop 2) consisted of only 66 individuals and, likely due to small sample size, produced poor quality allele assignments with high levels of homoplasy when analyzed by itself (data not shown). We therefore tested our method on Pop 1 (183 individuals) and on the combined set of 249 individuals.

For five out of eight loci, our algorithm found allele assignments devoid of homoplasy when only Pop 1 was used for assignment and when the allele swapping algorithm was used (Table 3). Eliminating the allele swapping algorithm or using the whole dataset for allele assignment increased the number of apparent homoplasious alleles in most cases, and did not reduce the number of apparent homoplasious alleles for any locus (Table 3). For the three loci with homoplasy, most genotypes in the dataset could not be assigned unambiguously (Table 3). For the five loci with no apparent homoplasy, nearly all genotypes in Pop 1 could be assigned unambiguously, and approximately one half to three quarters of the genotypes in Pop 2 (which was not used for creating the assignments) could be assigned unambiguously (Table 3). Despite the fact that Pop 1 was previously determined to consist of three subpopulations with pairwise Phi-PT [an *F_ST_* analog that can be used on both dominant and codominant markers (Peakall *et al*, 1995)] values ranging from 0.06 to 0.17 (Drauch Schreier *et al*, 2012), allelic variable correlations resulting from population structure did not appear to prevent us from obtaining reasonable allele assignments for the five loci without homoplasy. Significant positive correlations between allelic variables were found at one and two out of eight loci when Pop 1 and the whole dataset were used to make assignments, respectively (data not shown).

**Table 3:**
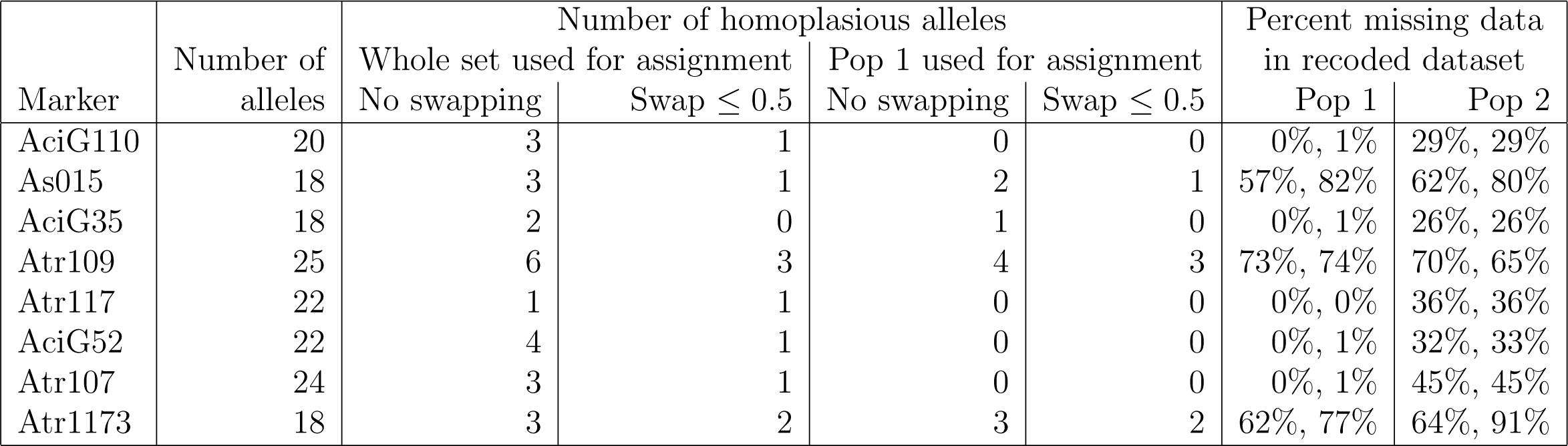
Assignment of alleles from eight microsatellite markers to two tetrasomic genomes in octoploid white sturgeon (*Acipenser transmontanus*). Alleles were assigned using negative correlations, with the exception of Atr117 in Pop 1 due to a fixed allele in that locus and population. Assignments were performed without allele swapping (“No swapping”, `rare.al.check = 0` in `testAlGroups`) and with allele swapping (“Swap ≤ 0:5”, `rare.al.check = 0.5`). In testing for homoplasy `testAlGroups` was run with the defaults of `tolerance = 0.05` to allow for 5% of genotypes to disagree with allele assignments, and `null.weight=0.5` to allow for the possibility of null alleles. Assignments were performed using the whole dataset of 249 individuals (“whole set”) or a subset of 183 individuals based on population structure (“Pop 1”, Supplementary Table 3 and Supplementary Fig. 1). The assignments from Pop 1 with Swap ≤ 0:5 were then used to split the dataset into isoloci using the `recodeAllopoly` function. Genotypes that could not be unambiguously determined were coded as missing data; percentages of missing data in each of two isoloci in Pop 1 and Pop 2 are shown.

| Marker | Number of alleles | Number of homoplasious alleles |  |  |  | Percent missing data in recoded dataset |  |
| --- | --- | --- | --- | --- | --- | --- | --- |
|  |  | Whole set used for assignment |  | Pop 1 used for assignment |  | Pop 1 | Pop 2 |
| | | No swapping | Swap $\leq 0.5$ | No swapping | Swap $\leq 0.5$ | | |
| AciG110 | 20 | 3 | 1 | 0 | 0 | 0%, 1% | 29%, 29% |
| As015 | 18 | 3 | 1 | 2 | 1 | 57%, 82% | 62%, 80% |
| AciG35 | 18 | 2 | 0 | 1 | 0 | 0%, 1% | 26%, 26% |
| Atr109 | 25 | 6 | 3 | 4 | 3 | 73%, 74% | 70%, 65% |
| Atr117 | 22 | 1 | 1 | 0 | 0 | 0%, 0% | 36%, 36% |
| AciG52 | 22 | 4 | 1 | 0 | 0 | 0%, 1% | 32%, 33% |
| Atr107 | 24 | 3 | 1 | 0 | 0 | 0%, 1% | 45%, 45% |
| Atr1173 | 18 | 3 | 2 | 3 | 2 | 62%, 77% | 64%, 91% |

By recoding allo-octoploid markers as tetrasomic isoloci, we were able to estimate allele frequencies, which would not have been possible otherwise. We were then able to use allele frequencies to estimate pairwise *G_ST_* between white sturgeon sampling regions. *G_ST_* estimates using recoded isoloci were very similar to estimates obtained using known tetrasomic microsatellite markers (Supplementary Fig. 2), suggesting that allele assignments were accurate. Out of the ten recoded isoloci, only one (Atr117_1) was consistently an outlier in terms of *G_ST_* estimates, giving especially high estimates between sampling regions corresponding to Pop 1 and Pop 2 (Supplementary Fig. 2, Supplementary Table 3). Atr117_1 had especially low genotype variability due to an allele that was present in all Pop 1 genotypes (Table 3), which likely accounted for the unusual *G_ST_* estimates at that isolocus. Otherwise, *G_ST_* estimates appeared unaffected by the large amounts of missing data introduced into Pop 2 by our method (Table 3, Supplementary Fig. 2), suggesting any bias in allele frequencies caused by the missing data was negligible.

## Simulated mapping populations

Negative correlations between allelic variables at the same isolocus can also occur in certain types of mapping populations, enabling the use of our algorithm to assign alleles to isoloci in these populations. There are several requirements that must be met however. 1) To prevent correlations between unlinked allelic variables, all individiduals in the population must be equally related to each other. Pedigrees, nested association mapping (NAM) populations, and multiple-cross mating designs are therefore not appropriate. 2) No allele should be present in all individuals in the population. Our method therefore cannot be used on backcross or near isogenic line (NIL) populations, which are expected to segregate only AB and BB genotypes. 3) All alleles belonging to one isolocus should have had the opportunity to pair with each other at meiosis. This eliminates F1 populations, where an individual with genotype AB might be crossed to an individual with genotype CD. However, allele assignments in F_2_ populations, as well as related populations such as recombinant inbred line (RIL) and doubled haploid (DH), can be peformed with very high accuracy using our algorithm.

Accuracy of allele assignment was 100% for allotetraploids and allohexaploids for all population types tested (F_2_ to F_8_; Table 4). Due to the highly heterozygous nature of tetrasomic loci, accuracy was 14% for allo-octoploids in the F_2_ generation. However, accuracy for allo-octoploids increased to 91% in the F_3_ and 100% in F_4_ and higher populations, due to increased homozygosity from selfing.

**Table 4:** Accuracy of allele assignment in mapping populations. Percentages of datasets with accurate allele assigments are shown. 95% confidence intervals are indicated. 1000 loci were simulated, each with 200 individuals.

| Generation | Allotetraploid | Allohexaploid | Allo-octoploid |
| --- | --- | --- | --- |
| F<sub>2</sub> | 100% ± 0% | 100% ± 0% | 13.6% ± 2.1% |
| F<sub>3</sub> | 100% ± 0% | 100% ± 0% | 91.4% ± 1.7% |
| F<sub>4</sub> | 100% ± 0% | 100% ± 0% | 100% ± 0% |
| F<sub>5</sub> | 100% ± 0% | 100% ± 0% | 100% ± 0% |
| F<sub>6</sub> | 100% ± 0% | 100% ± 0% | 100% ± 0% |
| F<sub>7</sub> | 100% ± 0% | 100% ± 0% | 100% ± 0% |
| F<sub>8</sub> | 100% ± 0% | 100% ± 0% | 100% ± 0% |

## Discussion

Here we introduce the R package polysat version 1.5, with several new functions applicable to the analysis of allopolyploids and diploidized autopolyploids. These include `simAllopoly`, which generates simulated datasets; `catalanAlleles`, which uses the the Catalán *et al* (2006) method to assign alleles to isoloci; `alleleCorrelations`, which performs Fisher’s exact test between each pair of allelic variables from a marker, and then uses K-means clustering and UPGMA to make initial assignments of alleles to isoloci; `testAlGroups`, which checks the consistency of allele assignments with individual genotypes, chooses between the K-means and UPGMA method, swaps alleles to different isoloci if it improves consistency, and identifies homoplasious alleles; `mergeAlleleAssignments`, which merges the allele assignments from two different populations using the same microsatellite marker; `processDatasetAllo`, which runs `alleleCorrelations`, `testAlGroups` (with multiple parameter sets), and `mergeAlleleAssignments` on an entire dataset; and `recodeAllopoly`, which uses allele assignments to recode the dataset, splitting each microsatellite marker into multiple isoloci. An overview of the data analysis workflow is given in Fig. 2. Previous versions of polysat (1.3 and earlier) were restricted in that estimation of allele frequency and certain inter-individual distance metrics could only be performed on autopolyploids. With the ability to assign alleles to isoloci, these parameters may now be estimated for allopolyploids as well.

We found that, with simulated data, the accuracy of our allele assignment algorithm was impacted by issues such as homoplasy and null alleles, and that the optimal parameters for the algorithm depended on which of these issues were present in the dataset. This suggests, since most users will not know whether their dataset has homoplasy or null alleles, that the `testAlGroups` function should initially be run with several different parameter sets, and for each locus, the results with the fewest homoplasious alleles should be chosen. A heatmap of the P-values generated from Fisher’s exact test can also serve as a qualitative visual indicator of how well the alleles can be separated into isolocus groups. We also found that, although our allele assignment algorithm was negatively impacted by meiotic error (pairing of non-homologous chromosomes during meiosis) and moderate population structure, its accuracy remained fairly high in both cases. Assuming correct allele assignments in a population with meiotic error, `recodeAllopoly` is able to identify some but not all individuals with meiotic error, for example if alleles A, B, and C belonged to one isolocus and D to another, an ABC D individual would be correctly recoded, where as an ABB D individual would be incorrectly recoded as AB DD. Otherwise, `recodeAllopoly` should give 100% accurate results if allele assignments are correct. Sensitivity to population structure is the biggest drawback of our method in comparison to that of Catalán *et al* (2006), which actually has improved results as population structure increases. However, even low frequencies of null alleles, homoplasy, or meiotic error can cause the method of Catalán *et al* (2006) to fail completely.

When discussing homoplasy with respect to our algorithm, we have referred exclusively to homoplasy between alleles belonging to different isoloci. It is important to note that homoplasy between alleles within an isolocus is also possible, meaning that two or more alleles belonging to one isolocus are identical in amplicon size but not identical by descent. Although such homoplasy is an important consideration for analyses that determine similarity between individuals and populations, homoplasy within isoloci does not affect the allele assignment methods described in this manuscript. Additionally, when discussing null alleles, we have assumed that non-null alleles still exist for all isoloci. It is also possible for an entire isolocus to be null. This is often apparent when a marker has fewer alleles per genotype than expected, *e.g.* a maximum of two alleles per individual in a tetraploid. Such loci should be excluded from the allele assignment analysis described in this manuscript. If they are included in an analysis accidentally, they can be identified by weak K-means/UPGMA clustering of alleles (which can be evaluated from the graphical output of `processDatasetAllo`) and by a high proportion of alleles appearing to be homoplasious.

Using a real microsatellite dataset from natural populations of white sturgeon, we found that our method was useful for recoding over half of the markers into two independently segregating isoloci each. Given that white sturgeon are octoploid with two tetrasomic subgenomes (Drauch Schreier *et al*, 2011), we expected this dataset to be problematic; having tetrasomic isoloci as opposed to disomic isoloci would reduce the magnitude of the negative correlations between allelic variables, and was observed in simulations to reduce the accuracy of assignment using our method, although not nearly as severly as the reduction in efficacy of the Catalán *et al* (2006) method (Supplementary Table 1, Fig. 3). In population genetic studies, we expect that microsatellite markers that can be recoded using our method could be used for analyses requiring polysomic or disomic inheritance [for example, estimation of allele frequency and population differentiation (Supplementary Fig. 2), Structure (Falush *et al*, 2007), or tests of Hardy-Weinberg Equilibrium], while the remaining markers will still be useful for other analysis (for example, Mantel tests using simple dissimilarity statistics). Additionally, we found that the allele assignments that we made were still moderately useful for recoding genotypes in a population that was not used for making the assignments. Despite the introduction of missing data into Pop 2 when its genotypes were recoded, *G_ST_* estimates were similar to those obtained from non-recoded tetrasomic microsatellites in the same populations (Supplementary Fig. 2). We do however recommend caution when interpreting results from loci where our method has introduced missing data for a large portion of individuals. Such results can be confirmed by comparison to results from loci with little or no missing data.

Although inappropriate for biallelic marker systems such as single nucleotide polymorphisms (SNPs) and dominant marker systems such as AFLPs, the method that we have described could theoretically be used to assign alleles to isoloci in any marker system in which multiple alleles are the norm. Allozymes, although rarely used in modern studies, are one such system. Although data from genotyping-by-sequencing (GBS, and the related technique restriction site-associated DNA sequencing, or RAD-seq) are typically processed to yield biallelic SNP markers, in the future as typical DNA sequencing read lengths increase, it may become common to find multiple SNPs within the physical distance covered by one read. In that case, haplotypes may be treated as alleles, and negative correlations between haplotypes may be used to assign them to isoloci.

## Obtaining polysat 1.5

To obtain polysat, first install the most recent version of R (available at http://www.r-project.org), launch R, then at the prompt type:

~~~
install.packages(”polysat”)
~~~

In the “doc” subdirectory of the package installation, PDF tutorials are available for polysat as a whole and for the methodology described in this manuscript. Source code is available at https://github.com/lvclark/polysat/ under a GNU GPL-2 license.

## Acknowledgements

Author LC was supported by the DOE Office of Science, Office of Biological and Environmental Research (grant number DE-SC0012379). We thank Subject Editor Frederic Austerlitz and three anonymous reviewers for feedback on an earlier version of this manuscript.

## Supporting Information

- polysatsupplementary.pdf: Supplementary materials and methods, tables, and figures.
- allopolyVignette.pdf: Tutorial for creating and using allele assignments in polysat.
- tables figs.R, sturgeontest.R: R scripts for reproducing the analyses in this manuscript.
- sturgeon.txt: White sturgeon microsatellite dataset used in strugeontest.R.

## Data Accessibility

polysat is available from CRAN (http://cran.r-project.org). All datasets and scripts used in this manuscript are provided as Supporting Information.

## Author Contributions

LC wrote and tested the software and drafted the manuscript. AS provided the white sturgeon data and gave critical feedback on the manuscript.

